# LncCE: Landscape of Cellular-Elevated LncRNAs in Single Cells Across Normal and Cancer Tissues

**DOI:** 10.1101/2024.05.17.594684

**Authors:** Kang Xu, Yujie Liu, Chongwen Lv, Ya Luo, Jingyi Shi, Haozhe Zou, Weiwei Zhou, Dezhong Lv, Changbo Yang, Yongsheng Li, Juan Xu

## Abstract

Long non-coding RNAs (lncRNAs) have emerged as significant players for maintaining the morphology and function of tissues or cells. The precise regulatory effectiveness of lncRNA is closely associated with the spatial expression patterns across tissues and cells. Here, we proposed the Cellular-Elevated LncRNA (LncCE) database to systematically explore cellular-elevated (CE) lncRNAs across normal and cancer tissues in single cells. LncCE encompasses 87,946 CE lncRNAs of 149 cell types by analyzing 181 single-cell RNA sequencing (scRNA-seq) datasets, involved in 20 fetal normal tissues, 59 adult normal tissues, as well as 32 adult and 5 pediatric cancer tissues.Two main search options were provided via a given lncRNA name or a cell type. The output results emphasize both qualitative and quantitative expression features of lncRNAs across different cell types, co-expression with protein-coding genes as well as their involved in biological functions. For cancers, LncCE particularly provided quantitative figures for exhibiting their expression changes compared to control samples and clinical associations with patient overall survivals. Together, LncCE offers an extensive, quantitative and user-friendly interface to investigate cellular-elevated expression atlas for lncRNAs across normal and cancers tissues at single-cell level. The LncCE database is available at http://bio-bigdata.hrbmu.edu.cn/LncCE.

## Introduction

Long non-coding RNAs (lncRNAs) are a class of noncoding RNAs which are longer than 200 nucleotides (nt) and lack protein coding capacity [1-3]. It was widely accepted that lncRNAs were closely associated with various crucial biological functions to maintain tissue morphology, even cancer development [4-6].

Emerging studies have comprehensively characterized the expression patterns of genes with bulk transcriptomes. Besides tissue-specific (TS) genes, other two kinds of genes were found to be important for the realization of the physiological function of normal tissues as well as the formation and development of cancer, including tissue-enhanced genes and tissue enriched ones [7, 8]. These three categories of genes all exhibit tissue-elevated expression in a certain tissue, that is, the gene expression is dominantly higher in some tissues than in other tissues. Genes with tissue-elevated expression patterns were constantly identified and validated across normal and cancer tissues [9-11], and their biomarker potential for cancer diagnosis and prognosis were discussed [6, 12]. Thus, comprehensive characterization of spatial expression can reveal the function of lncRNAs across tissues and cancers. Indeed, current developments in transcriptome analyses unveiled the stronger tissue-specificity expression of lncRNAs than protein-coding genes [13]. In our previous studies, the tissue-elevated expression patterns of lncRNAs have been systematically explored across normal and cancer tissues, and their key roles were also revealed in maintaining morphology and function of tissues [6, 12].

Single-cell RNA sequencing (scRNA-seq) technologies are powerful for analyzing the spatial expression patterns of genes at single cell level [14], providing an unprecedented chance to highlight increasingly challenging biological questions and explore the molecular mechanisms related to carcinogenesis [15-17]. Emerging studies also have investigated the expression pattern of lncRNA across normal and cancer cells [18-21], and further experimentally validated lncRNAs with cell state-specific functions involved in cell cycle progression and apoptosis [22, 23]. For example, lncRNAs specially expressed in T cell were found to play important roles in cancer immunity [24]. However, yet little is known about lncRNA expression properties at the single-cell level, and there is still no comprehensive database for providing CE lncRNA expression atlas across human tissues and cancers via large-scale scRNA-seq data.

To address these challenges, we constructed the public resource, LncCE (http://bio-bigdata.hrbmu.edu.cn/LncCE), which is a landscape of Cellular-Elevated lncRNAs in single-cell across normal and cancer tissues. The cellular-elevated (CE) lncRNAs were comprehensively discovered, which were further classified into three categories based on their dominant in a certain cell type, including cell specific (CS), cell enriched (CER) and cell enhanced (CEH). LncCE not only provides the cellular-elevated expression patterns of lncRNAs, but also can be used for downstream analysis, such as identifying novel cellular markers and comparing cellular-elevated expression patterns of lncRNAs across different cellular states. LncCE could significantly help the research community to understand the biological functions of lncRNA in cells, tissues and tumorigenesis.

## Data collection and processing

### Data collection and pre-processing in LncCE

For single-cell RNA transcriptome resources across normal tissues, three widely available transcriptome datasets were collected (**Table S1**), including Human Cell Landscape (HCL) [25], Cross-tissue Immune Cell Atlas (TICA) [26], The Tabula Sapiens (TTS) [27]. Specially, one normal fetal transcriptome datasets, HCL Fetal datasets, also had been contained in our study for a comprehensive compare of CE (cellular-elevated) lncRNAs of development.

For single-cell RNA transcriptome resources across cancer tissues, we collected scRNA raw count files from Gene Expression Omnibus (GEO) [28], ArrayExpress [29], TISCH [30] and TISCH2 [31]. Similarly, pediatric cancer transcriptome datasets were also collected, including five pediatric cancer types [32, 33].

For all scRNA transcriptome resources, we also collected the metadata information (including sample ID, organ/tissue origin, clinical treatment, biosample groups, and major cell types) of each normal or cancer tissue. For all scRNA datasets, we removed genes that were not expressed in at least 3 cells and cells that did not have at least 50 detected genes. The data was then normalized using scale factor of 10,000 and natural log-transformed, and then LncCE documented 2,893,787 cells from 181 transcriptome resources, including 74 transcriptomes of 32 cancers, 5 transcriptomes of 5 pediatric cancers, 82 transcriptomes of 59 normal tissues and 20 transcriptomes of 20 fetal normal tissues. In the cell type identification, LncCE used the original major cell type annotation from the metadata information. To provide comprehensive and precise annotations of similar cell types, we have corrected and unified the cell types, such as, in some datasets the cell names are misspelled or the same cell type have different names in different datasets. Furthermore, to identify clusters of distinct cell populations, the tSNE clustering algorithms was implemented in the Seurat R package (v4.3.0).

### Gene annotation

Gene annotation files were downloaded from GENCODE (release 38, GRCh38) [34] which includes different types of genes, including protein-coding genes, long noncoding RNAs (lncRNAs), and pseudogenes. For a more comprehensive analysis of lncRNAs expression patterns at single-cell level, we consider ‘pseudogenes’ as ‘long noncoding RNAs’ in our study.

### Identifying the CE lncRNA

To identify CE lncRNAs in each tissue, we identified lncRNAs that have at least 5-fold higher expression levels in one cell-type compared with all other cell-types [6, 9, 12, 35]. Moreover, CE lncRNAs were also further classified into three subcategories to reflect increasing degrees of elevated expression in a particular tissue, including “cell specific (CS)”, “cell enriched (CER)”, and “cell enhanced (CEH)”. (i) CS lncRNAs were expressed only in a particular cell-type, where the expression thresholds were set 0.001 for counts per million (CPM); (ii) CER lncRNAs were with at least 5-fold higher expression level in a particular cell-type compared with the max expression levels in all other cell-types; and (iii) CEH lncRNAs were with at least 5-fold higher expression level in a particular cell-type compared with the average expression levels in all other cell-types. Furthermore, we defined a CE lncRNA when it expressed in more than ten cells of the same cell-types. Similarly, we also identified cancer CE lncRNAs in each cancer cell-type. Moreover, CE protein-coding genes in each cell-type under normal or cancer states were also identified.

### LncRNA-mRNA correlation

Due to the inefficiency of the technical molecule, scRNA-Seq may not be able to detect the truly expressed gene in some cells and is therefore represented by false zero expression. Thus, scRNA-seq data may be much sparser than the whole tissue RNA sequencing, we apply scLink, a new method to better characterize the statistical dependencies between genes in single cell [36]. Users can set different correlation thresholds, spatial pattern of mRNAs to visualize the co-expressed lncRNA-mRNA subnetwork based on the tool echarts. Moreover, CE mRNAs in the same cell were also highlighted in the co-expression subnetwork by node colors.

### Function prediction of CE lncRNAs

Function prediction of CE lncRNAs also provided in single-cell. After users select the co-expressed mRNAs in the above step, genes are online subjected into the R packages “clusterProfiler” to predict enriched functions of lncRNA, including Gene Ontology categories and pathways [37].

### Database construction

LncCE was constructed by Java Server Pages and deployed on Tomcat software (v6). All datasets in LncCE were documented and managed in MySQL database (v5.5). Several commonly used Java script packages, including ECharts (v5.2.2), Datatable (v1.12.1), Highcharts (v7.1.2) and plotly (v2.16.1), were implemented for presentation of query results and interactive visualization of data. All data processing and integration analysis were performed using R software (v4.1.2). Currently, the website has been tested on several popular web browsers, including Google Chrome (preferred), Firefox, or Apple Safari browsers.

## Database content and usage

### Data summary

The current version of LncCE includes 2,893,787 cells, covering 149 cell types across 37 cancer types and 79 normal tissues of the human body. There were 20 fetal normal tissues, 59 adult normal tissues, 32 adult and 5 pediatric cancer tissues. On average, each scRNA-seq dataset has 34,865 cells, ranging from 515 to 103,703 cells (**Table S1**). A total of 14,941 lncRNAs were identified in LncCE database. The CE lncRNAs were comprehensively identified (details in data collection and processing), which were further classified into three categories, including CS lncRNAs, CER lncRNAs and CEH lncRNAs. Currently, there were 87,946 CE lncRNAs were curated in LncCE (**Table S2**), with the largest number of CE lncRNAs in lacrimal gland functional unit cell of eye (TTS, n = 3342) and the smallest number of CE lncRNAs in B cell of BRCA (GSE114727_indrop, n = 1).

### Database overview

LncCE is not only a comprehensive resource of CE lncRNAs across single-cell but also provides a user-friendly web interface for investing the spatial expressions of lncRNA across cellular states (**Figure 1**). The users can switch between adult and pediatric normal/cancer tissues by typing the button on the homepage. Users could click on the corresponding button in the homepage to enter the “Browse”, “Search”, and “Download” pages for browsing, searching, and downloading all CE lncRNAs in LncCE.

**Figure 1.**
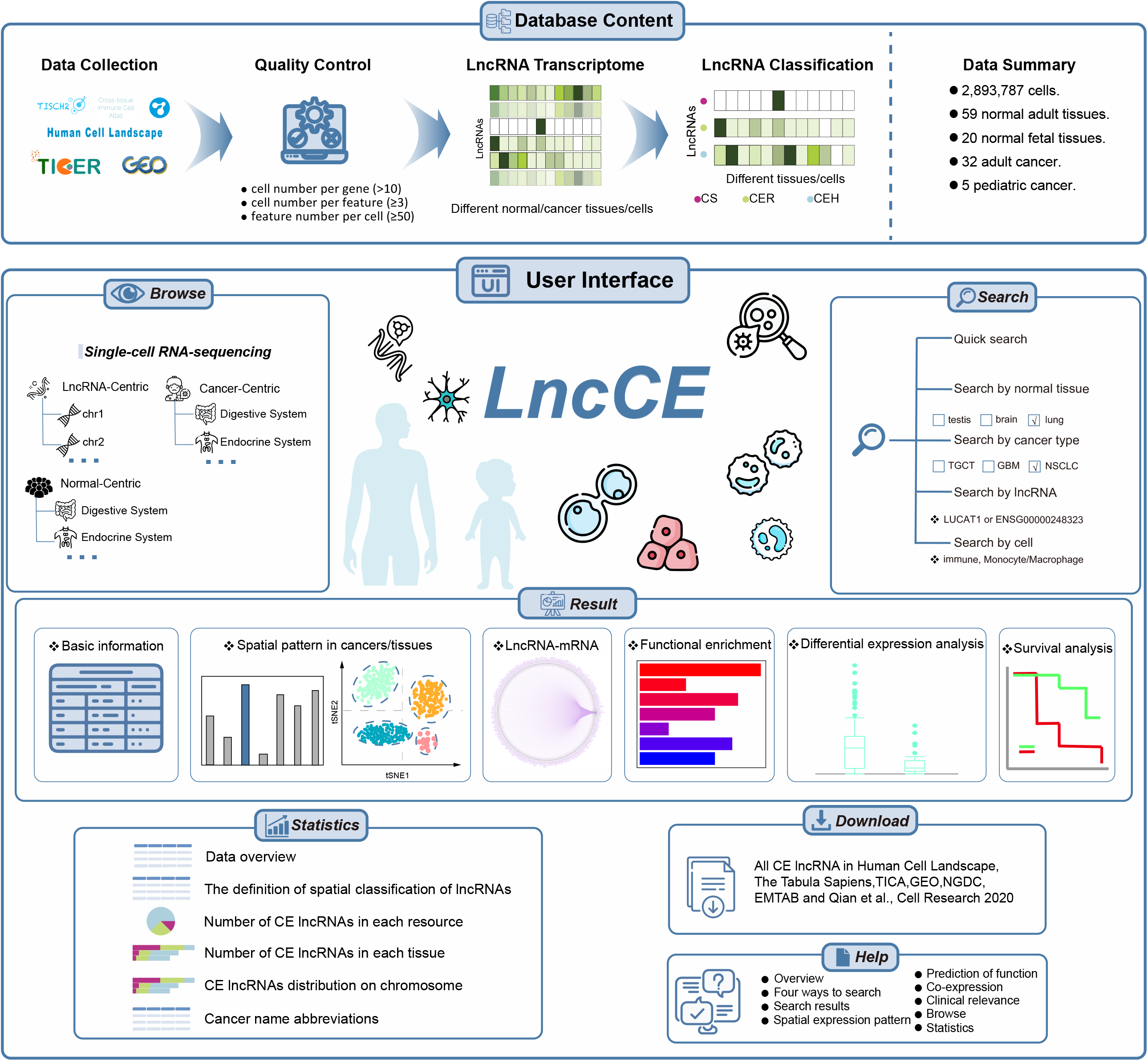
Schematic of overall design of LncCE. Top, workflow of obtaining all CE lncRNAs from multiple scRNA-seq datasets. Middle and bottom, user interface of LncCE. The users can select adult or fetal normal tissues, or adult or pediatric cancer tissues for quick queries. In this panel, the Search, Browse, Statistics, Download, and Help modules provide flexible ways to access the dataset. Four types of search moudles were provides for all CE lncRNAs. Three browsing ways are also given.

CE lncRNAs could be browsed by “LncRNA-Centric”, “Normal-Centric” and “Cancer-Centric”. In LncRNA-Centric page, all CE lncRNAs were organized in the hierarchical structure based on chromosomal localization. In the other two browse pages, normal, adult, and pediatric cancer tissues were also organized in hierarchical structure based on anatomic classification in human body map. Moreover, users can quickly enter the “Searching Result” pages by clicking tissue of interest in the human body on the home page. In the “Search” sections, four different query options were provided on the basis of normal or cancer tissues of interest, lncRNA names or cell types. In addition, the statistic information of LncCE can also be accessed from the “Statistics” page. All data in the database can be freely downloaded from the “Download” page. A detailed tutorial showing how to browse and query data was also available on the “Help” page.

### LncRNA-based exploration with LncCE

Recently, some studies have reported that *LUCAT1* (lung cancer associated transcript 1) is a cancer-related and myeloid cell-specific lncRNA, which can interact with *STAT1* (signal transducer and activator of transcription 1) to inhibit ISGs (interferon-stimulated genes) transcription [38-40]. As use case for a “LncRNA” search, we employed LncCE to identify the cell types expressing *LUCAT1* across different tissues.

Upon entering a lncRNA in the corresponding search field at the “LncRNA” region of the “search” page, the LncCE displays a table of the gene’s basic information in each cell type in every dataset in which the gene is identified as CE lncRNA (**Figure 2A and B**). We searched for *LUCAT1* and observed that it is most highly expressed in myeloid cells in most cancers including lung cancer and colorectal cancer, and also highly expressed in myeloid cells in normal tissues including lung, esophagus tissues (**Figure 2A**).

**Figure 2.**
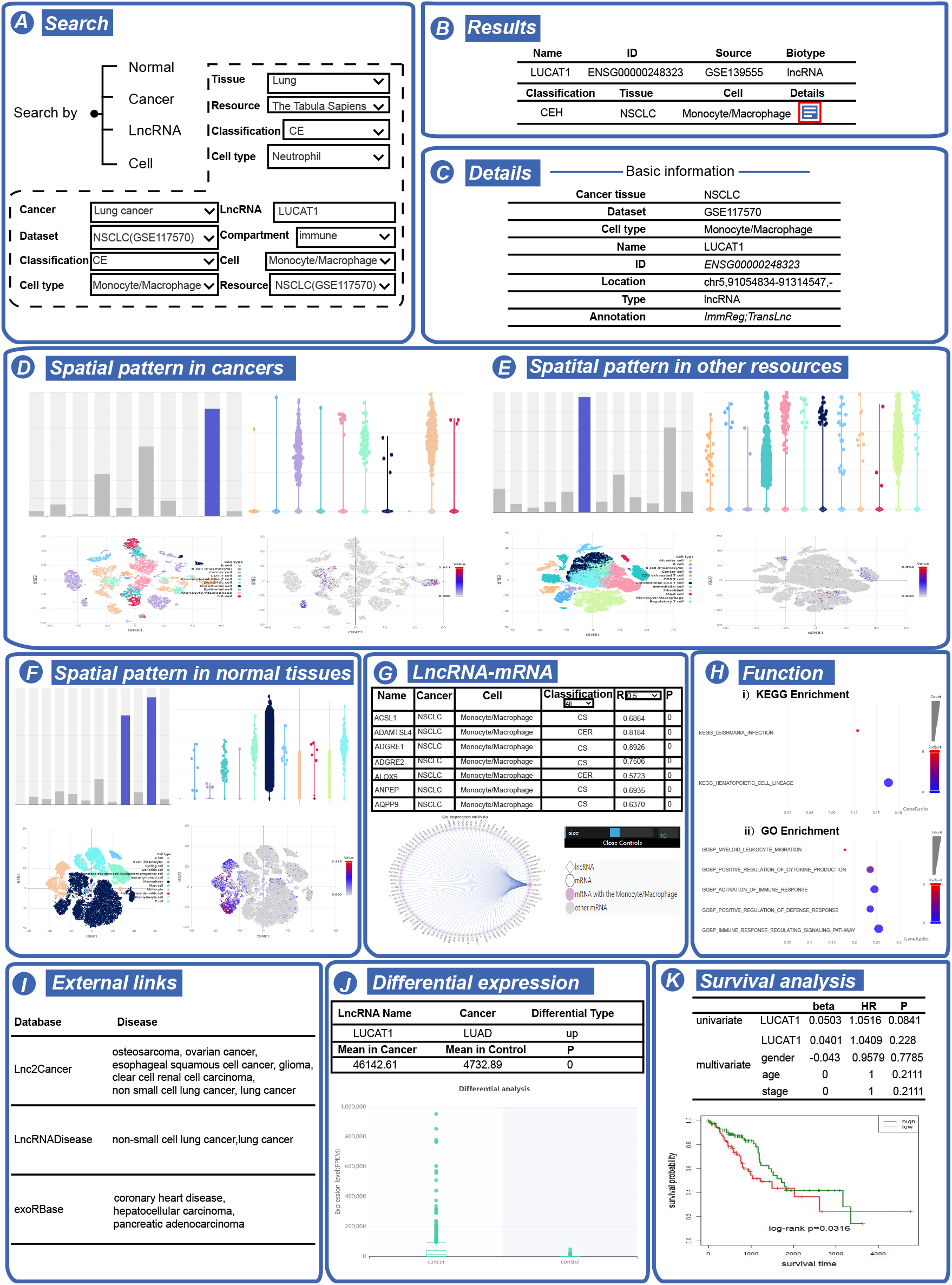
LncRNA-based exploration with LncCE. **A**. Search by adult or fetal normal tissues, or adult or pediatric cancer tissues, lncRNAs or cell types of interests. **B**. The result list for lncRNAs. **C**. Basic annotation information for cellular-elevated lncRNA *LUCAT1* and annotations in other relevant databases. **D**. A global map of different cell populations and expression levels of *LUCAT1* across cell types in CE cancer tissue (NSCLC) in the selected dataset (GSE117570). **E**. A global map of different cell populations and expression levels of *LUCAT1* across cell types in CE cancer tissue (NSCLC) in the other dataset. **F**. A global map of different cell populations and expression levels of *LUCAT1* across cell types in normal tissues (Lung) associated with CE cancer tissue (NSCLC). **G**. Co-expression network between CE lncRNA *LUCAT1* and mRNAs. **H**. Functional and pathway enrichment analysis of co-expression mRNAs. **I**. External links related to NSCLC and *LUCAT1*. **J**. Expression and regulation of lncRNA *LUCAT1* in TCGA cancer. **K**. Survival analysis of CE lncRNA *LUCAT1* in TCGA cancer.

By clicking the details button in these tables, users can further obtain more details for individual entry. A hyperlink was linked to the detail result page for the CE lncRNA *LUCAT1* in NSCLC (non-small cell lung cancer) of GSE117570. Eight major types of information were provided (**Figure 2C-2K**). (i) Basic annotation information was provided and an annotation link could provide annotations of *LUCAT1* in ImmReg [41], TransLnc [42] and LncSpA [6]. (ii) CE cancer tissue (NSCLC), subclassification of CE, and corresponding expression levels were listed in a table, a bar chart and box chart of expression across cell types was provided, and a tSNE figure which is colored by cell type was used to visualize the expression of lncRNA, such as *LUCAT1*. (iii) The qualitative and quantitative spatial expression patterns in normal tissues (lung tissue as its CE tissue) were provided. (iv) Co-expression between CE lncRNA *LUCAT1* and mRNAs were shown in a network view. In addition, the correlation information was listed in a table, and users could select different thresholds (0-0.7) to filter the lncRNA-mRNA co-expression network. (v) Co-expressed mRNAs were used for functional and pathway enrichment analysis, identifying various kinds of relations with physiologic and pathologic lung tissue. (vi) Evidences have also shown the association between NSCLC and *LUCAT1* from Lnc2Cancer, LncRNADisease, and exoRBase. (vii) Expressions of lncRNA in cancer. We found that *LUCAT1* was upregulated in LUAD (lung adenocarcinoma) and LUSC (lung squamous cell carcinoma) in TCGA, suggesting *LUCAT1* as a candidate oncogenic lncRNA, which is consistent with the study of Agarwal S. et.al [38]. (viii) The results of regression analysis and the Kaplan–Meier survival plot indicated that *LUCAT1* was a protective factor in LUAD and LUSC. Taken together, these eight panels provided detailed information for understanding the function of CE lncRNA across different cell types under normal and cancer tissues.

### Cell type-based exploration with LncCE

Emerging studies have well-characterized the function of cell-type elevated mRNAs across normal and cancer tissues, but there has been less research to lncRNAs. LncCE represents a comprehensive annotation of cell-type elevated lncRNAs, providing the possibility to investigate the function of CE lncRNAs. LncRNAs are cell-type specifically expressed in a variety of cell types (**Table S3**), including immune cells (T cell, B cell, macrophage and etc.), stromal cell, endothelial cell, and muscle cell.

Next, we focused on CE lncRNAs which were identified in no less than 15 datasets (**Figure 3**). Several lncRNAs, including *LUCAT1, MIAT* (myocardial infarction associated transcript), *WFDC21P* (WAP four-disulfide core domain 21), *CARMN* (cardiac mesoderm enhancer-associated non-coding RNA) and *PCAT19* (prostate cancer associated transcript 19), appear to be expressed in matched cell types, suggesting a conserved role in these cell types. *LUCAT1* is reported as a myeloid-specific lncRNA, and it is identified as CE lncRNA in 85 datasets in LncCE (**Figure 4A**), in particular with myeloid dervied cells (macrophage, monocyte and neutrophil, 66/85) and myeloid cell (7/85). *MIAT* is a T cell marker in LncCE which is identified as CE lncRNA in 20 adult cancer datasets (**Figure 4B**), and it has been reported mainly expressed in tumor and T cells which indicating that *MIAT* may be involved in the immune escape process of cancer [43]. *WFDC21P*, also known as *lnc-DC*, is a lncRNA expressed exclusively in human dendritic cells, that is required for optimal dendritic cell differentiation from human monocytes and *WFDC21P* has been shown to promotes the nuclear translocation and function of *STAT3* by interacting with the transcription factors *STAT3* [44], and is identified as CE lncRNA of dendritic cell in 23 datasets (**Figure S1A**). *CARMN* has been reported an evolutionarily conserved smooth muscle cell-specific lncRNA [45] and is identified as CE lncRNA in endothelial and muscle cells (**Figure S1B**). *PCAT19* has been reported to safeguard DNA in quiescent endothelial cells by preventing uncontrolled phosphorylation of *RPA2* (replication protein A2) [46] and drive prostate cancer [47] or as a prognostic biomarker for endometrial cancer [48]. In LncCE, *PCAT19* is identified as CE lncRNA in 126 datasets, particularly in endothelial cell (108/126), suggesting *PCAT19* may be a novel biomarker for endothelial cell (**Figure S1C**).

**Figure 3.**
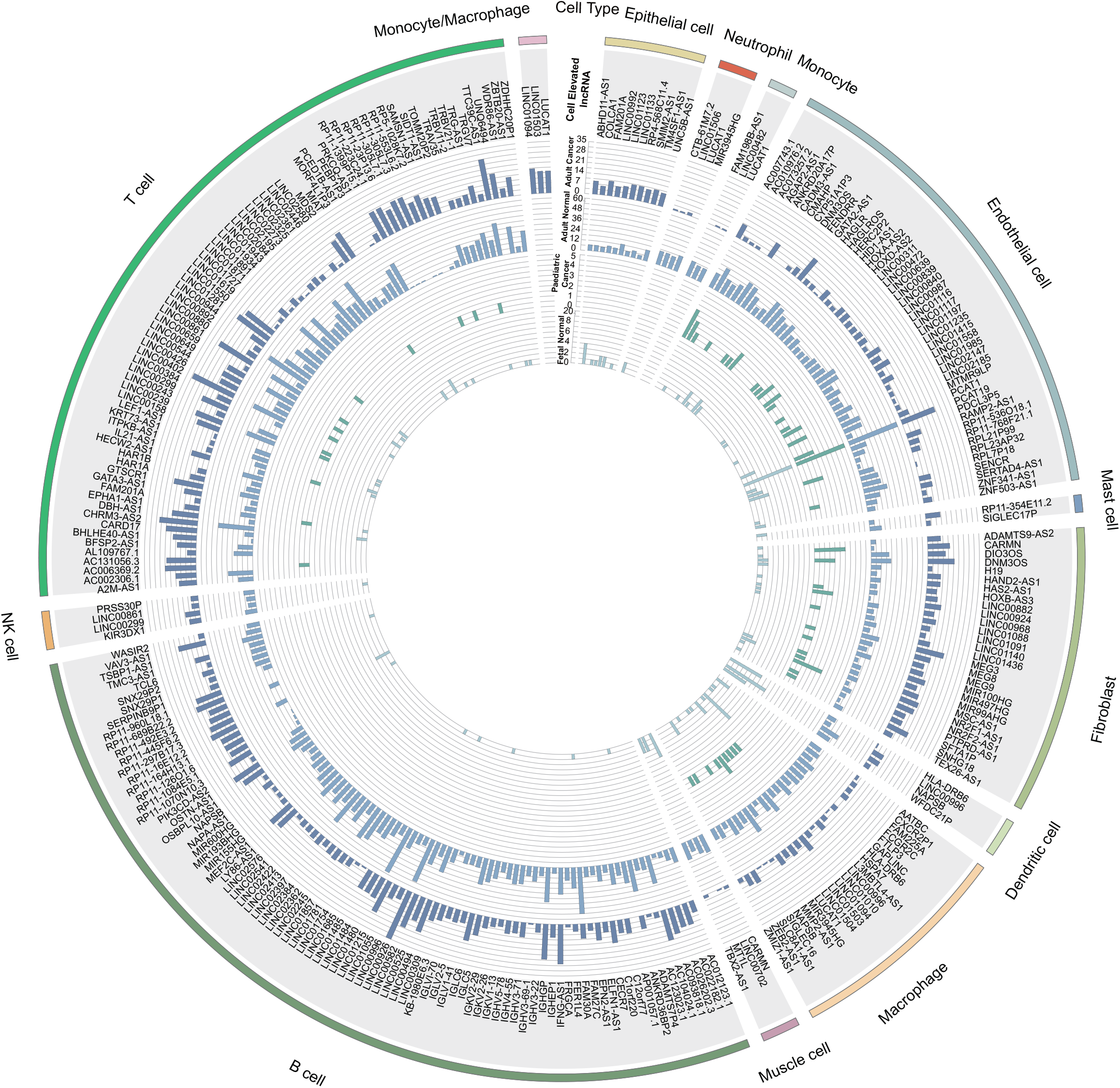
Distribution of CE lncRNA in each cell type (n ≥ 15) across adult and fetal normal tissues, adult and pediatric cancer tissues. The outermost circle is the cell type, the length shows the amount of CE lncRNAs. The penultimate circle is CE lncRNA names. The third last circle is the distribution of CE lncRNAs count in adult cancer tissues, followed by adult normal tissues, pediatric cancer tissues and fetal normal tissues. The height of the bar shows the number of datasets that the lncRNA is identified as CE lncRNA.

**Figure 4.**
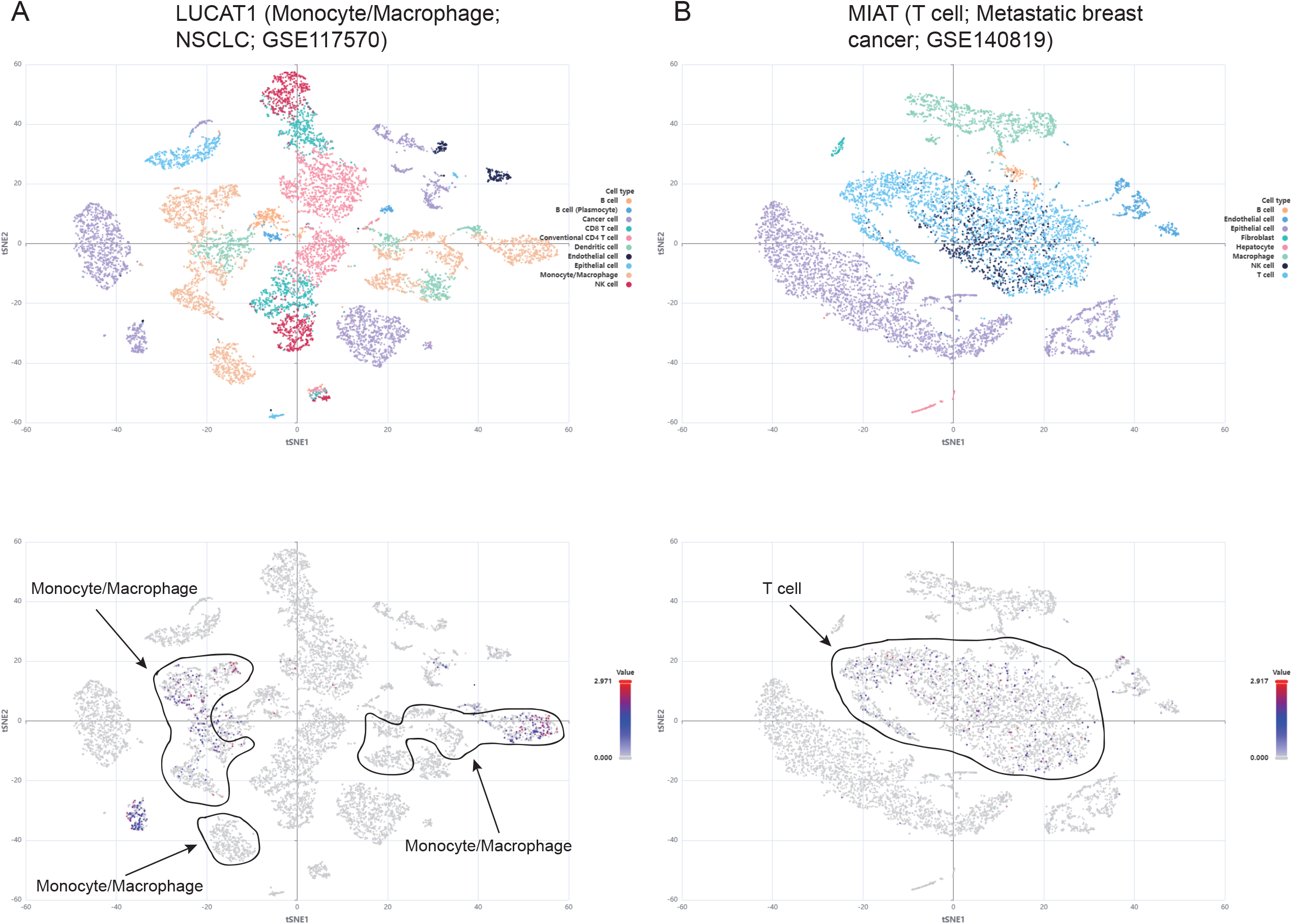
Cell type-based exploration with LncCE. **A**. tSNE plots of GSE117570_NSCLC dataset colored by cell type and *LUCAT1* expression. B. tSNE plots of GSE140819_Metastatic_breast_cancer dataset colored by cell type and *MIAT* expression.

These observations suggested a considerable number of cell elevated lncRNAs are involved in cellular differentiation, activation and inflammation based signaling, cancer initiation, development and treatment resistance.

## Discussion

In summary, LncCE is a comprehensive resource for investigating the spatial expression patterns of lncRNAs at single-cell level across adult and fetal normal tissues, as well as adult and pediatric cancer types. User-friendly interface was designed for querying, browsing, and downloading the CE lncRNAs of interest. Eight major types of information for CE lncRNAs were provided for visualizing and understanding their function in physiologic and pathologic phenotypes.

Comparing with the other resources, LncCE is particularly dedicated to cellular-elevated lncRNA across normal and cancer single-cell transcriptomic datasets (**Table S4**). LncCE not only provides the spatial expression patterns of CE lncRNAs, but also can be used for downstream analysis, such as identifying novel cellular markers and comparing spatial patterns of lncRNAs across different states. In the future, we will continue to update LncCE to include more cells across normal and cancer tissues and maintain it as a valuable resource. In addition, the drug susceptibility for CE lncRNAs will also be added to our database in the future. CE lncRNAs in LncCE are potentially promising candidate therapeutic targets in precision oncology. The CE lncRNAs may be the marker genes for distinguishing cell types. We believe that LncCE will be a valuable resource for both experimental and computational researchers to bridge the knowledge gap from lncRNA expression to phenotypes.

## Data availability

The online database LncCE is publicly available at http://bio-bigdata.hrbmu.edu.cn/LncCE.

## CRediT author statement

**Kang Xu:** Formal Analysis, Data curation, Writing - original draft, Visualization. **Yujie Liu:** Software, Writing - original draft, Visualization. **Chongwen Lv:** Data curation, Writing - original draft, Visualization. **Ya Luo:** Data curation, Visualization. **Jingyi Shi:** Resources, Visualization. **Haozhe Zou:** Data Curation, Resources. **Weiwei Zhou:** Data Curation, Resources. **Dezhong Lv:** Software, Data Curation. **Changbo Yang:** Visualization. **Yongsheng Li:** Conceptualization, Funding acquisition, Writing - review & editing, Supervision. **Juan Xu:** Conceptualization, Funding acquisition, Writing - review & editing, Supervision. All authors have read and approved the final manuscript.

## Acknowledgments

This research was funded by National Natural Science Foundation of China [32170676, 31970646, 32060152 and 32070673]; Natural Science Foundation of Heilongjiang Province (Key Program) [ZD2023C007]; the China Brain Project [2021ZD0202403], and Heilongjiang Touyan Innovation Team Program.

## Supplementary material

**Figure S1.**
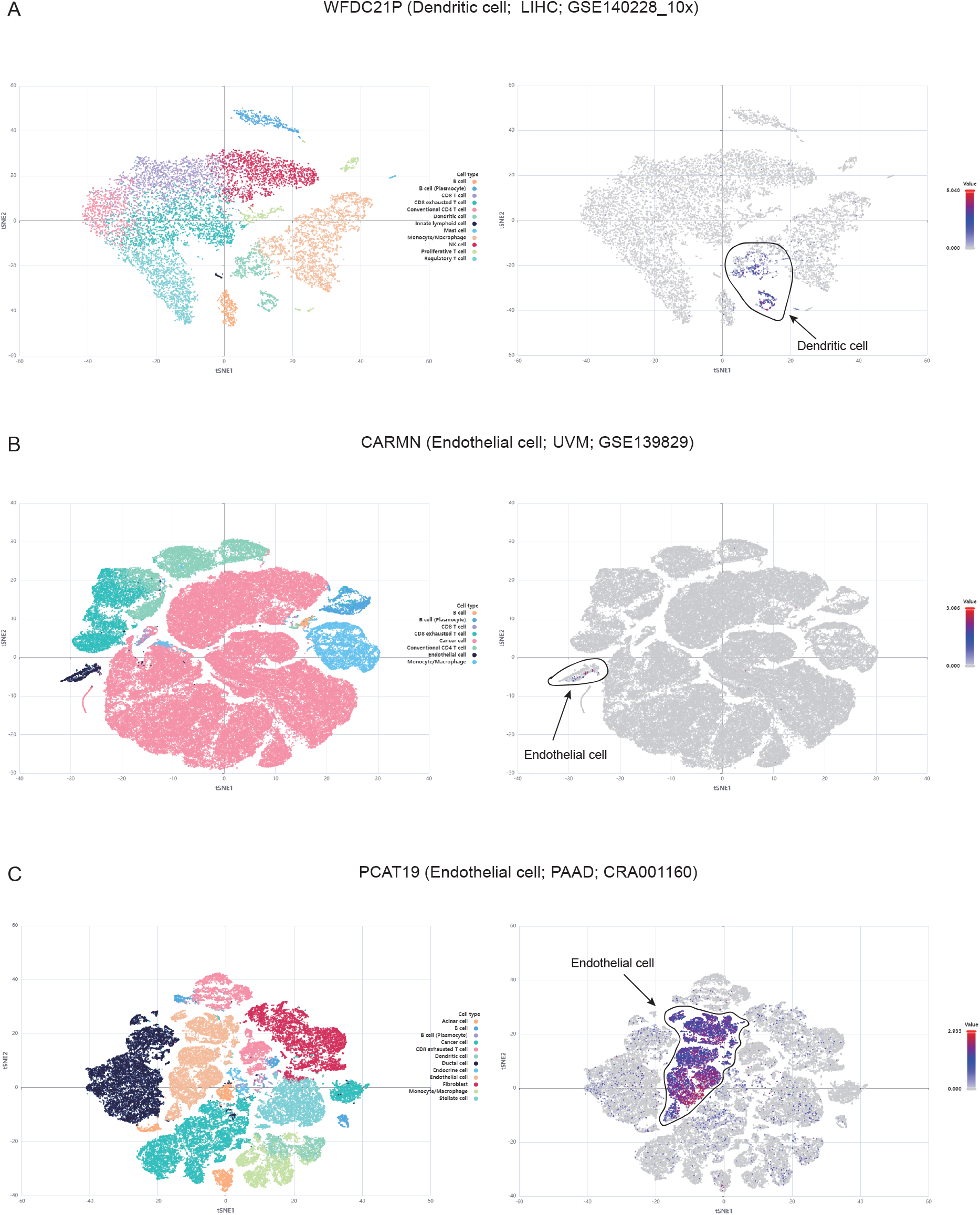
tSNE plots of GSE140228_LIHC (A), GSE139829_UVM (B) and CRA001160_PAAD (C) dataset colored by cell type and CE lncRNAs (*WFDC21P, CARMN* and *PCAT19*) expression

**Table S1 The information of all datasets**

**Table S2 The number of CE lncRNA of each datasets across adult/pediatric cancer/normal tissues**

**Table S3 The list of CE lncRNAs for each cell type**

**Table S4 Highlights of lncCE comparing with other human scRNA-seq database**

## References

[1] Kapranov P, Cheng J, Dike S, Nix DA, Duttagupta R, Willingham AT, et al. RNA maps reveal new RNA classes and a possible function for pervasive transcription. Science 2007;316:1484–8.

[2] Wang KC, Chang HY. Molecular mechanisms of long noncoding RNAs. Mol Cell 2011;43:904–14.

[3] Sarropoulos I, Marin R, Cardoso-Moreira M, Kaessmann H. Developmental dynamics of lncRNAs across mammalian organs and species. Nature 2019;571:510–4.

[4] Li Y, Li L, Wang Z, Pan T, Sahni N, Jin X, et al. LncMAP: Pan-cancer atlas of long noncoding RNA-mediated transcriptional network perturbations. Nucleic Acids Res 2018;46:1113–23.

[5] Li Y, Jiang T, Zhou W, Li J, Li X, Wang Q, et al. Pan-cancer characterization of immune-related lncRNAs identifies potential oncogenic biomarkers. Nat Commun 2020;11:1000.

[6] Lv D, Xu K, Jin X, Li J, Shi Y, Zhang M, et al. LncSpA: LncRNA Spatial Atlas of Expression across Normal and Cancer Tissues. Cancer Res 2020;80:2067–71.

[7] Fagerberg L, Hallstrom BM, Oksvold P, Kampf C, Djureinovic D, Odeberg J, et al. Analysis of the human tissue-specific expression by genome-wide integration of transcriptomics and antibody-based proteomics. Mol Cell Proteomics 2014;13:397–406.

[8] Uhlen M, Fagerberg L, Hallstrom BM, Lindskog C, Oksvold P, Mardinoglu A, et al. Proteomics. Tissue-based map of the human proteome. Science 2015;347:1260419.

[9] Uhlen M, Hallstrom BM, Lindskog C, Mardinoglu A, Ponten F, Nielsen J. Transcriptomics resources of human tissues and organs. Mol Syst Biol 2016;12:862.

[10] Shi Y, Chen L, Liotta LA, Wan HH, Rodgers GP. Glia maturation factor gamma (GMFG): a cytokine-responsive protein during hematopoietic lineage development and its functional genomics analysis. Genomics Proteomics Bioinformatics 2006;4:145–55.

[11] Amaral PP, Mattick JS. Noncoding RNA in development. Mamm Genome 2008;19:454–92.

[12] Xu K, Jin X, Luo Y, Zou H, Lv D, Wang L, et al. Spatial transcriptome analysis of long non-coding RNAs reveals tissue specificity and functional roles in cancer. J Zhejiang Univ Sci B 2023;24:15–31.

[13] Rinn JL, Chang HY. Genome Regulation by Long Noncoding RNAs. Annual Review of Biochemistry, Vol 81 2012;81:145–66.

[14] Ziegenhain C, Vieth B, Parekh S, Reinius B, Guillaumet-Adkins A, Smets M, et al. Comparative Analysis of Single-Cell RNA Sequencing Methods. Mol Cell 2017;65:631–43 e4.

[15] Ziaee S, Boroumand MA, Salehi R, Sadeghian S, Hosseindokht M, Sharifi M. Non-invasive diagnosis of early-onset coronary artery disease based on cell type-specific gene expression analyses. Biomed Pharmacother 2018;108:1115–22.

[16] Saviano A, Henderson NC, Baumert TF. Single-cell genomics and spatial transcriptomics: Discovery of novel cell states and cellular interactions in liver physiology and disease biology. J Hepatol 2020;73:1219–30.

[17] Joanito I, Wirapati P, Zhao N, Nawaz Z, Yeo G, Lee F, et al. Single-cell and bulk transcriptome sequencing identifies two epithelial tumor cell states and refines the consensus molecular classification of colorectal cancer. Nat Genet 2022;54:963–75.

[18] Sanmarco LM, Rone JM, Polonio CM, Fernandez Lahore G, Giovannoni F, Ferrara K, et al. Lactate limits CNS autoimmunity by stabilizing HIF-1alpha in dendritic cells. Nature 2023;620:881–9.

[19] Fyfe I. Single-cell atlas maps cell-specific gene changes in Alzheimer disease. Nat Rev Neurol 2020;16:1.

[20] Ner-Gaon H, Melchior A, Golan N, Ben-Haim Y, Shay T. JingleBells: A Repository of Immune-Related Single-Cell RNA-Sequencing Datasets. J Immunol 2017;198:3375–9.

[21] Cao Y, Zhu J, Jia P, Zhao Z. scRNASeqDB: A Database for RNA-Seq Based Gene Expression Profiles in Human Single Cells. Genes (Basel) 2017;8.

[22] Johnsson P, Ziegenhain C, Hartmanis L, Hendriks GJ, Hagemann-Jensen M, Reinius B, et al. Transcriptional kinetics and molecular functions of long noncoding RNAs. Nat Genet 2022;54:306–17.

[23] Kim DH, Marinov GK, Pepke S, Singer ZS, He P, Williams B, et al. Single-cell transcriptome analysis reveals dynamic changes in lncRNA expression during reprogramming. Cell Stem Cell 2015;16:88–101.

[24] Luo H, Bu D, Shao L, Li Y, Sun L, Wang C, et al. Single-cell Long Non-coding RNA Landscape of T Cells in Human Cancer Immunity. Genomics Proteomics Bioinformatics 2021;19:377–93.

[25] Han X, Zhou Z, Fei L, Sun H, Wang R, Chen Y, et al. Construction of a human cell landscape at single-cell level. Nature 2020;581:303–9.

[26] Dominguez Conde C, Xu C, Jarvis LB, Rainbow DB, Wells SB, Gomes T, et al. Cross-tissue immune cell analysis reveals tissue-specific features in humans. Science 2022;376:eabl5197.

[27] Tabula Sapiens C, Jones RC, Karkanias J, Krasnow MA, Pisco AO, Quake SR, et al. The Tabula Sapiens: A multiple-organ, single-cell transcriptomic atlas of humans. Science 2022;376:eabl4896.

[28] Barrett T, Wilhite SE, Ledoux P, Evangelista C, Kim IF, Tomashevsky M, et al. NCBI GEO: archive for functional genomics data sets--update. Nucleic Acids Res 2013;41:D991–5.

[29] Athar A, Fullgrabe A, George N, Iqbal H, Huerta L, Ali A, et al. ArrayExpress update - from bulk to single-cell expression data. Nucleic Acids Res 2019;47:D711–D5.

[30] Sun D, Wang J, Han Y, Dong X, Ge J, Zheng R, et al. TISCH: a comprehensive web resource enabling interactive single-cell transcriptome visualization of tumor microenvironment. Nucleic Acids Res 2021;49:D1420–D30.

[31] Han Y, Wang Y, Dong X, Sun D, Liu Z, Yue J, et al. TISCH2: expanded datasets and new tools for single-cell transcriptome analyses of the tumor microenvironment. Nucleic Acids Res 2023;51:D1425–D31.

[32] Slyper M, Porter CBM, Ashenberg O, Waldman J, Drokhlyansky E, Wakiro I, et al. A single-cell and single-nucleus RNA-Seq toolbox for fresh and frozen human tumors. Nat Med 2020;26:792–802.

[33] Gillen AE, Riemondy KA, Amani V, Griesinger AM, Gilani A, Venkataraman S, et al. Single-Cell RNA Sequencing of Childhood Ependymoma Reveals Neoplastic Cell Subpopulations That Impact Molecular Classification and Etiology. Cell Rep 2020;32:108023.

[34] Frankish A, Diekhans M, Jungreis I, Lagarde J, Loveland JE, Mudge JM, et al. Gencode 2021. Nucleic Acids Res 2021;49:D916–D23.

[35] Uhlen M, Zhang C, Lee S, Sjostedt E, Fagerberg L, Bidkhori G, et al. A pathology atlas of the human cancer transcriptome. Science 2017;357.

[36] Vivian Li W, Li Y. scLink: Inferring Sparse Gene Co-expression Networks from Single-cell Expression Data. Genomics Proteomics Bioinformatics 2021;19:475–92.

[37] Wu T, Hu E, Xu S, Chen M, Guo P, Dai Z, et al. clusterProfiler 4.0: A universal enrichment tool for interpreting omics data. Innovation (Camb) 2021;2:100141.

[38] Agarwal S, Vierbuchen T, Ghosh S, Chan J, Jiang Z, Kandasamy RK, et al. The long non-coding RNA LUCAT1 is a negative feedback regulator of interferon responses in humans. Nat Commun 2020;11:6348.

[39] Sun Y, Jin SD, Zhu Q, Han L, Feng J, Lu XY, et al. Long non-coding RNA LUCAT1 is associated with poor prognosis in human non-small lung cancer and regulates cell proliferation via epigenetically repressing p21 and p57 expression. Oncotarget 2017;8:28297–311.

[40] Luzon-Toro B, Fernandez RM, Martos-Martinez JM, Rubio-Manzanares-Dorado M, Antinolo G, Borrego S. LncRNA LUCAT1 as a novel prognostic biomarker for patients with papillary thyroid cancer. Sci Rep 2019;9:14374.

[41] Jiang TT, Zhou WW, Chang ZH, Zou HZ, Bai J, Sun QS, et al. ImmReg: the regulon atlas of immune-related pathways across cancer types. Nucleic Acids Research 2021;49:12106–18.

[42] Lv DZ, Chang ZH, Cai YY, Li JY, Wang LP, Jiang QS, et al. TransLnc: a comprehensive resource for translatable lncRNAs extends immunopeptidome. Nucleic Acids Research 2022;50:D413–D20.

[43] Peng L, Chen Y, Ou Q, Wang X, Tang N. LncRNA MIAT correlates with immune infiltrates and drug reactions in hepatocellular carcinoma. Int Immunopharmacol 2020;89:107071.

[44] Wang P, Xue Y, Han Y, Lin L, Wu C, Xu S, et al. The STAT3-binding long noncoding RNA lnc-DC controls human dendritic cell differentiation. Science 2014;344:310–3.

[45] Dong K, Shen J, He X, Hu G, Wang L, Osman I, et al. CARMN Is an Evolutionarily Conserved Smooth Muscle Cell-Specific LncRNA That Maintains Contractile Phenotype by Binding Myocardin. Circulation 2021;144:1856–75.

[46] Oo JA, Palfi K, Warwick T, Wittig I, Prieto-Garcia C, Matkovic V, et al. Long non-coding RNA PCAT19 safeguards DNA in quiescent endothelial cells by preventing uncontrolled phosphorylation of RPA2. Cell Rep 2022;41:111670.

[47] Hua JT, Ahmed M, Guo H, Zhang Y, Chen S, Soares F, et al. Risk SNP-Mediated Promoter-Enhancer Switching Drives Prostate Cancer through lncRNA PCAT19. Cell 2018;174:564–75 e18.

[48] Wang Z, Liu Y, Zhang J, Zhao R, Zhou X, Wang H. An Immune-Related Long Noncoding RNA Signature as a Prognostic Biomarker for Human Endometrial Cancer. J Oncol 2021;2021:9972454.

